# Actin filaments regulate microtubule growth at the centrosome

**DOI:** 10.1101/302190

**Authors:** Daisuke Inoue, Dorian Obino, Francesca Farina, Jérémie Gaillard, Christophe Guerin, Laurent Blanchoin, Ana-Maria Lennon-Duménil, Manuel Théry

## Abstract

The centrosome is the main microtubule-organizing centre. It also organizes a local network of actin filaments. However, the precise function of the actin network at the centrosome is not well understood. Here we show that increasing densities of actin filaments at the centrosome of lymphocytes were correlated with reduced amounts of microtubules. Furthermore, lymphocyte activation resulted in centrosomal-actin disassembly and an increase in microtubule number. To further investigate the direct crosstalk between actin and microtubules at the centrosome, we performed in vitro reconstitution assays based on (i) purified centrosomes and (ii) on the co-micropatterning of microtubule seeds and actin filaments. The two assays demonstrated that actin filaments perturb microtubule growth by steric hindrance. Finally, we showed that cell adhesion and spreading leads to lower densities of centrosomal actin thus resulting in higher microtubule growth. Hence we propose a novel mechanism by which the number of centrosomal microtubules is regulated by cell adhesion and actin-network architecture.

## Introduction

The growth of the microtubule network and its architecture regulates cell polarisation, migration and numerous key functions in differentiated cells (de Forges et al., 2012; Sanchez and Feldman, 2016; Mimori-Kiyosue, 2011; Etienne-Manneville, 2013). Microtubule growth first depends on microtubule nucleation, which is regulated by large complexes serving as microtubule templates and proteins that stabilise early protofilament arrangements (Roostalu and Surrey, 2017; Wieczorek et al., 2015). Then, microtubule elongation becomes regulated by microtubule-associated proteins and molecular motors acting at the growing end of microtubules (Akhmanova and Steinmetz, 2015). The architecture of the microtubule network - the spatial distribution and orientation of microtubules - is heavily influenced by its biochemical interactions and physical interplay with actin filaments (Rodriguez et al., 2003; Coles and Bradke, 2015; Huber et al., 2015). Although the physical crosslinking of the two networks can occur at any points along microtubule length,(Mohan and John, 2015) the sites of intensive crosstalk occur at the growing ends of microtubules (Akhmanova and Steinmetz, 2015).

The growth of microtubules can also be directed by actin-based structures (Théry et al., 2006; López et al., 2014; Kaverina et al., 1998). They can force the alignment of microtubules (Elie et al., 2015), resist their progression (Burnette et al., 2007), capture, bundle or stabilise them (Zhou et al., 2002; Hutchins and Wray, 2014), submit them to mechanical forces (Gupton et al., 2002; Fakhri et al., 2014; Robison et al., 2016) or define the limits in space into which they are confined (Katrukha et al., 2017). The actin-microtubule interplay mostly takes place at the cell periphery, because most actin filaments are nucleated at and reorganized into actin-based structures near the plasma membrane (Blanchoin et al., 2014). We recently have identified a subset of actin filaments that form at the centrosome at the cell centre (Farina et al., 2016). The centrosome is the main microtubule nucleating and organizing centre of the cell and sustains the highest concentration of microtubules in the cell. Centrosomal-actin filaments have been shown to be involved in several function including centrosome anchoring to the nucleus (Obino et al., 2016), centrosome separation in mitosis (Au et al., 2017) and ciliary-vesicle transport in the early stages of ciliogenesis (Wu et al., 2018). Whether centrosomal-actin filaments affect centrosomal microtubules is not yet known.

Here we investigated how the processes of actin and microtubule growth at the centrosome influence each other. We provide in vitro and in vivo evidence that these two processes are antagonistic, most likely as a result of physical hindrance. Our results further suggest that the antagonism from the centrosomal actin filaments restricts microtubule growth in response to cell adhesion.

## Results

### The centrosomal-actin network appears to negatively regulate the microtubule network in B lymphocytes

B-lymphocyte polarization can be achieved by B-cell receptor (BCR) activation from binding surface-tethered cognate antigens, and requires the local reduction of centrosomal actin density (Obino et al., 2016). To evaluate how microtubules were affected in resting and activated B lymphocytes, we examined by fluorescent microscopy of fixed cells, microtubule density throughout the cell in comparison with changes to the density of centrosomal actin filaments (Figure 1). As expected, B lymphocyte activation was associated with a lower density (by 30%) of actin at the centrosome (Obino et al., 2016). It was also associated with a higher density (by 20%) of microtubules (Figure 1B, C). A closer analysis by single cells showed a clear negative correlation between centrosomal-actin density and microtubules density in resting lymphocytes (r=–0.44) which appeared to be maintained but shifted toward lower actin densities and higher microtubule amount in activated lymphocytes (Figure 1D), suggesting that the interplay between the two networks is not specific to the activation but an intrinsic relationship.

**Figure 1:**
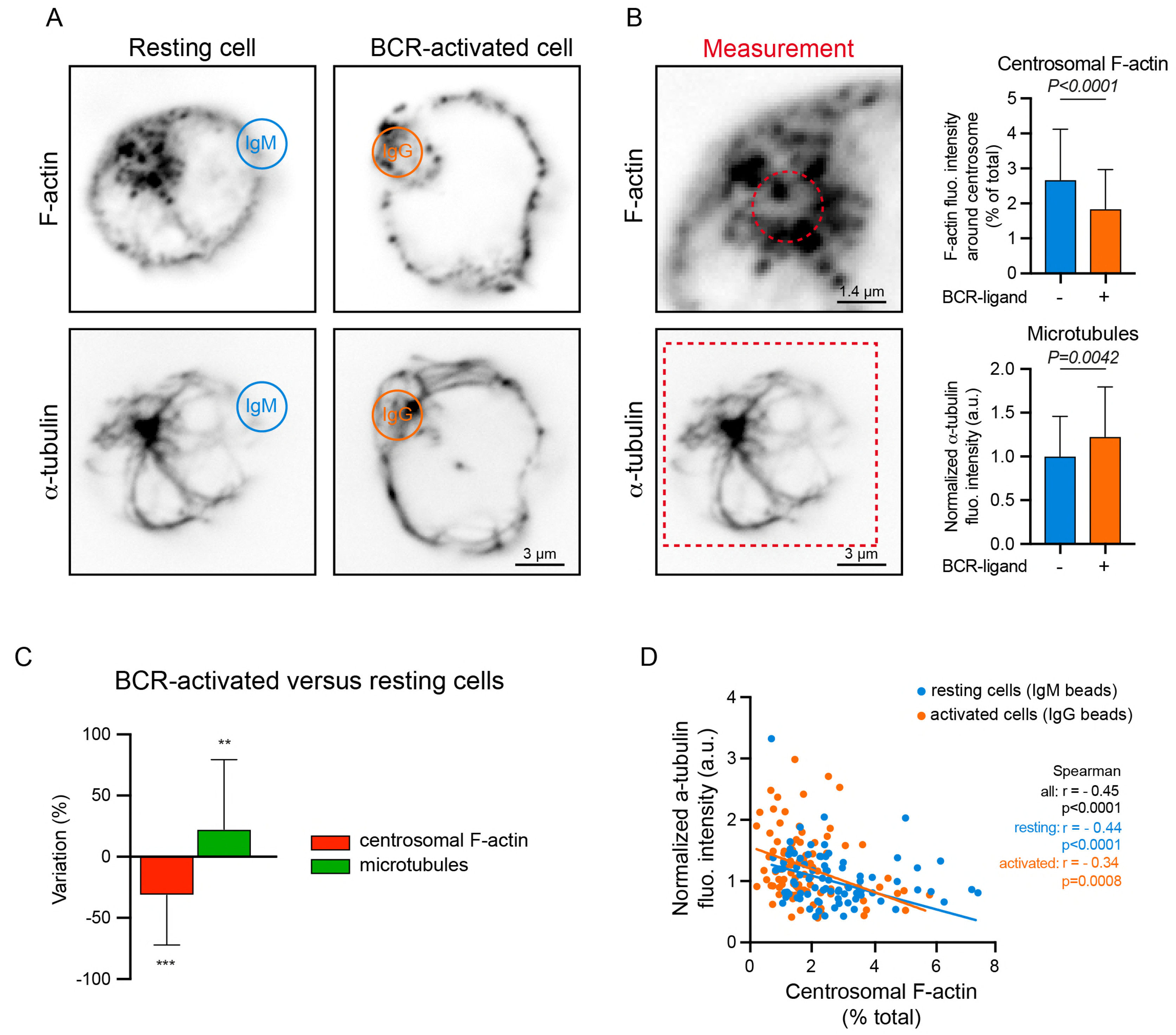
Cytoskeleton remodelling in B lymphocytes upon antigen stimulation. A- IIA1.6 B lymphoma cells were stimulated with BCR-ligand^−^ (IgM) or BCR-ligand^+^ (IgG) beads for 60 min, fixed and stained for and F-actin (top) and α-tubulin (bottom). Scale bar represents 3 μm. B- Histograms show the quantifications of the total amount of polymerized fluorescent tubulin (bottom, values were normalized with respect to the mean of control condition) and filamentous F-actin at the centrosome (top, dashed outline, values correspond to the fraction of fluorescence in a 1.6-micron-wide area around the centrosome relative to the total fluorescence in the cell). Measurements were pooled from 3 independent experiments; IgM (BCR-ligand^−^): n= 88; IgG (BCR-ligand^+^): n= 93. Error bars correspond to standard deviations. Scale bars represent 1.4 μm (top) and 3 μm (bottom). C- Percentage differences of centrosomal F-actin and microtubule fluorescence intensities in cells stimulated with BCR-ligand^+^ beads with respect to cells stimulated with BCR-ligand^−^ beads. Errors bars represent standard deviations. D- The graph shows the same measurements as panel B in an XY representation of individual measurements. The two lines correspond to linear regressions of the two sets of data relative to cells stimulated with BCR-ligand^+^ (activated cells) or BCR-ligand^−^ (resting cells) beads.

To test the hypothesis that the density of centrosomal actin is driving the reduction in microtubule density, B lymphocytes were treated with actin-filament inhibitors (Figure 2A). Treatment with the actin polymerization inhibitors (Arp2/3 inhibitor CK666) or latrunculin A reduced the centrosomal actin density and increased the microtubule density throughout the cell (Figure 2B, C), thus supporting the hypothesis. Conversely, treatment with the Formin inhibitor SMIFH2, increased centrosomal actin density and marginally decreased microtubule density throughout the cell (Figure 2B, C), thus confirming the negative relationship between the two networks. Overall, the analysis of individual cells showed a negative correlation between centrosomal actin filaments and microtubules. The inhibition of formin and Arp2/3 induced higher and lower actin densities at the centrosome, respectively, and thus expanded the range in which the negative correlation could be observed (Figure 2D).

**Figure 2:**
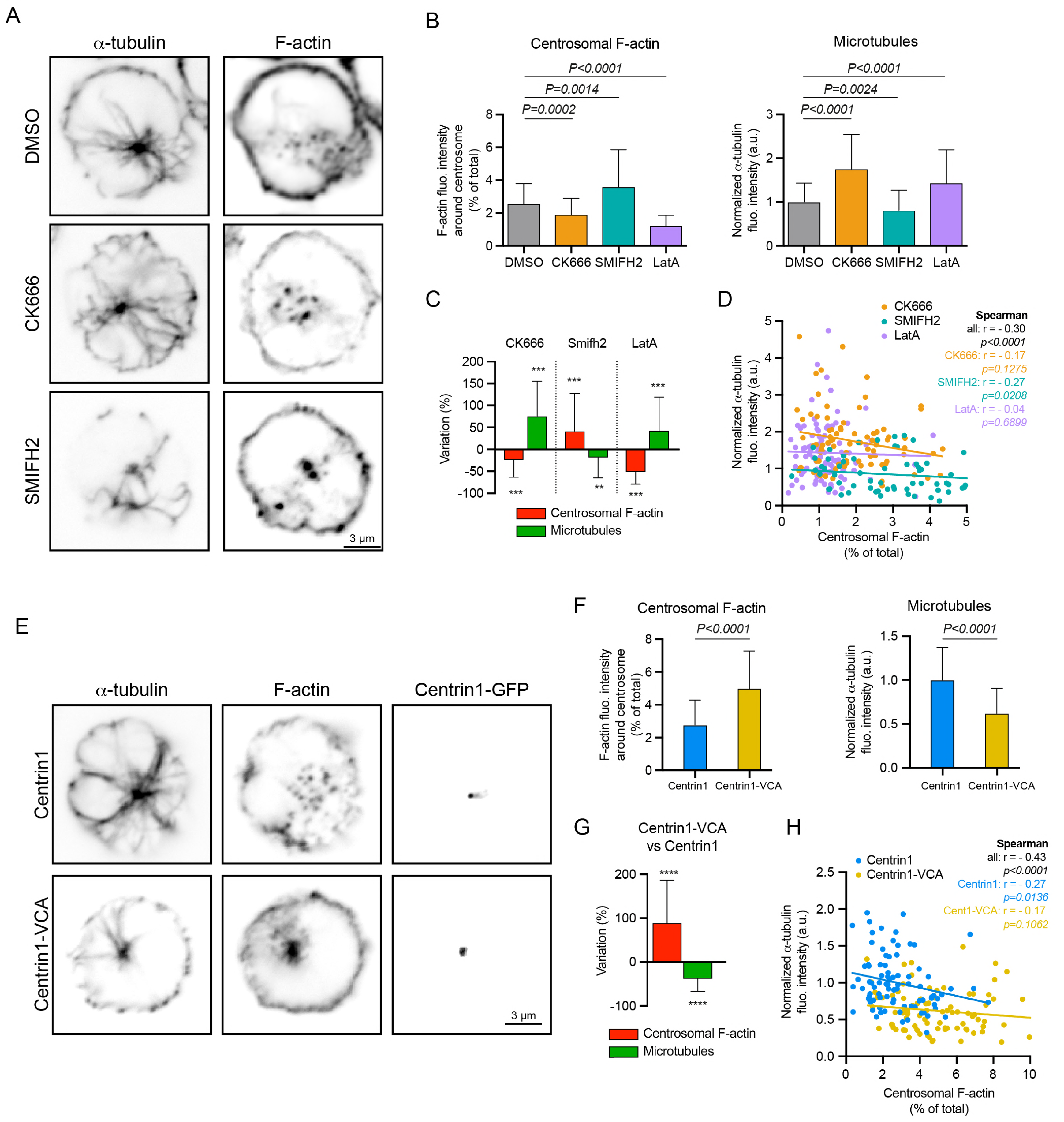
The impact of modulating centrosomal actin on microtubules in B lymphocytes. A- IIA1.6 B lymphoma cells were treated 45 min with indicated inhibitors (CK666 at 25 μM, SMIFH2 at 25μM; latrunculin-A 5μM) or DMSO as control prior to being fixed and stained for α-tubulin (left column) and F-actin (right column). Scale bar represents 3 μm. B- Histograms show the quantifications of the total amount of polymerized fluorescent tubulin (right, values were normalized with respect to the mean of control condition) and filamentous actin at the centrosome (left, values correspond to the fraction of fluorescence in a 1.6-micron-wide area around the centrosome relative to the total fluorescence in the cell). Measurements were pooled from 3 independent experiments; DMSO: n= 91, CK666: n= 82, SMIFH2: n= 74, latrunculinA: n= 96. Error bars correspond to standard deviations. C- Percentage differences of centrosomal F-actin and microtubule fluorescence intensities in cells treated with cytoskeleton inhibitors in comparison with the respective densities in cells treated with DMSO. Errors bars represent standard deviations. D- The graph shows the same measurements than in panel B in an XY representation of individual measurements. The three lines correspond to linear regressions of the three sets of data relative to cells stimulated with each actin drug. E- IIA1.6 B lymphoma cells were transfected to transiently express centrin1-VCA-GFP (bottom) or centrin1-GFP (top) as control prior to be fixed and stained for α-tubulin (left column) and F-actin (middle column). The GFP signal of centrin1 or centrin1-VCA is shown in the right column to illustrate the proper centrosome targeting. Scale bar represents 3 μm. F- Histograms show the quantifications of the total amount of polymerized fluorescent tubulin (right, values were normalized with respect to the mean of the control condition) and filamentous actin at the centrosome (left, values correspond to the fraction of fluorescence in a 1.6-micron-wide area around the centrosome relative to the total fluorescence in the cell). Measurements were pooled from 3 independent experiments; centrin1-GFP: n= 88, centrin1-VCA-GFP: n= 87. Errors bars represent standard deviations. G- Percentage differences of centrosomal F-actin and microtubule fluorescence intensities in cells transfected with centrin1-VCA-GFP in comparison with the respective densities in cells transfected with centrin1-GFP. Errors bars represent standard deviations. H- The graph shows the same measurements as in panel F in an XY representation of individual measurements. The two lines correspond to linear regressions of the two sets of data relative to cells transfected with centrin1-VCA-GFP or centrin1-GFP.

Given that the actin polymerization inhibitors would affect actin throughout the cell and not only at the centrosome, we next examined B-lymphocytes which expressed a fusion protein (centrin1-VCA-GFP; (Obino et al., 2016)) that promotes actin filament nucleation at the centrosome (Figure 2E). Hence the expression of centrin1-VCA-GFP strongly increased the density of centrosomal-actin filaments and decreased the microtubule density throughout the cell supporting the specific role of actin filaments at the centrosome in the negative regulation of the microtubule network (Figure 2F, G, H).

### The centrosomal-actin network perturbs the eloboration of the microtubule network in vitro

A limitation to the interpretation of the B-lymphocyte experiments was that local perturbations to the actin network could have affected other actin networks in the same cell by a process of actin-network homeostasis that operates throughout the cell (Burke et al., 2014; Suarez and Kovar, 2016; Suarez et al., 2014). Therefore, an increase in actin density at the centrosome could have been offset by a corresponding decrease in actin density elsewhere in the cell (e.g. in cytoplasmic and cortical networks). To circumvent this limitation, we used an in vitro model that reconstituted actin and microtubule networks from actin monomers and tubulin dimers incubated in the presence of a centrosome labelled with centrin1-GFP. In this model and as expected (Farina et al., 2016), 25% of the centrosomes (i.e. centrin1-GFP positive puncta) were associated with actin and microtubule networks (Figure 3A). Among those centrosomes, the actin density per centrosome was negatively correlated with the number of microtubules per centrosome (Figure 3B). Actin-filament density at the centrosome was then altered by incubating centrosomes in different concentrations of free actin monomers, with the tubulin monomer concentration kept constant (Figure 3C). Consistent with the hypothesis, higher actin concentrations were associated with lower microtubule numbers per chromosome (Figure 3D). Moreover, the highest actin concentration almost completely inhibited microtubule growth (Figure 3D). These results from the in vitro experiments suggest that actin filaments perturb microtubule formation at the centrosome. Therefore it is plausible that in the B lymphocyte experiments, the centrosomal-actin network had direct and antagonistic effects on the microtubule network emanating from the centrosome.

**Figure 3:**
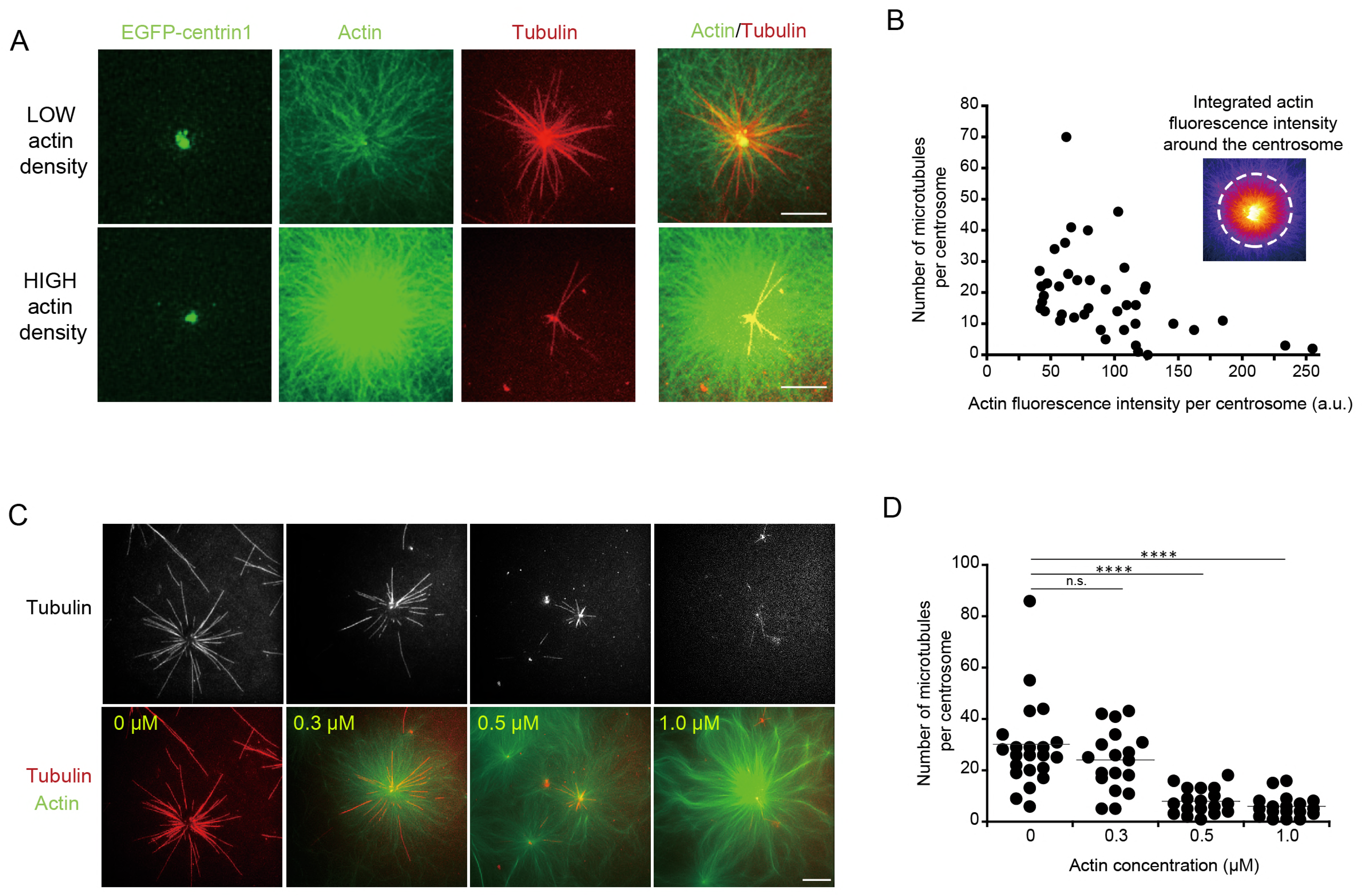
Assembly of microtubules and actin filaments on isolated centrosomes. A- Two sets of representative images showing fluorescent microtubules and actin filaments assembled from isolated centrosomes. Centrosomes were isolated from Jurkat cells expressing centrin1-GFP. Upper and lower lines show actin filaments and microtubules radiating from two distinct centrosomes with low (top) and high (bottom) densities of actin filaments. Scale bars represent 10 μm. B- The graph shows the number of microtubules per centrosome relative to the density of actin filaments. Inset shows actin filaments at the centrosome with a FIRE look-up table and a 20 μm-wide circle in which actin fluorescence intensity is measured. Measurements were pooled from 5 independent experiments. C- Microtubules (top line) and actin filaments (bottom line) assembly from isolated centrosomes in the presence of increasing concentration of monomeric actin (from left to right). Scale bar represents 20 μm. D- The graph shows the number of microtubules per centrosome in response to increasing concentrations of monomeric actin. Data were pooled from 2 independent experiments.

To further explore the dynamics of the interaction between the centrosomal-actin network and the microtubule network, the in vitro model was manipulated by sequential addition of the network components. By incubating with tubulin dimers first, microtubules formed in the absence of actin filaments (Figure 4A, B). When actin monomers were introduced (together with tubulin dimers to maintain the tubulin-dimer concentration), the number of microtubules increased on all centrosomes, irrespective of whether the centrosomes triggered the formation of actin filaments or not (Figure 4C). An explanation for this unexpected observation was that the addition of new tubulin dimers increased the effective concentration of free tubulins. Furthermore, not all centrosomes were capable of nucleating actin filaments, and there was no difference in the microtubule numbers per centrosome between those centrosomes with and those without actin filaments (Figure 4C). This suggested that in this model, the stability of preassembled microtubules may not be sensitive to actin filaments that form at the microtubule ends proximal to the centrosome, and newly assembled microtubules could form in spaces along pre-existing microtubules or in spaces created from depolymerized microtubules.

**Figure 4:**
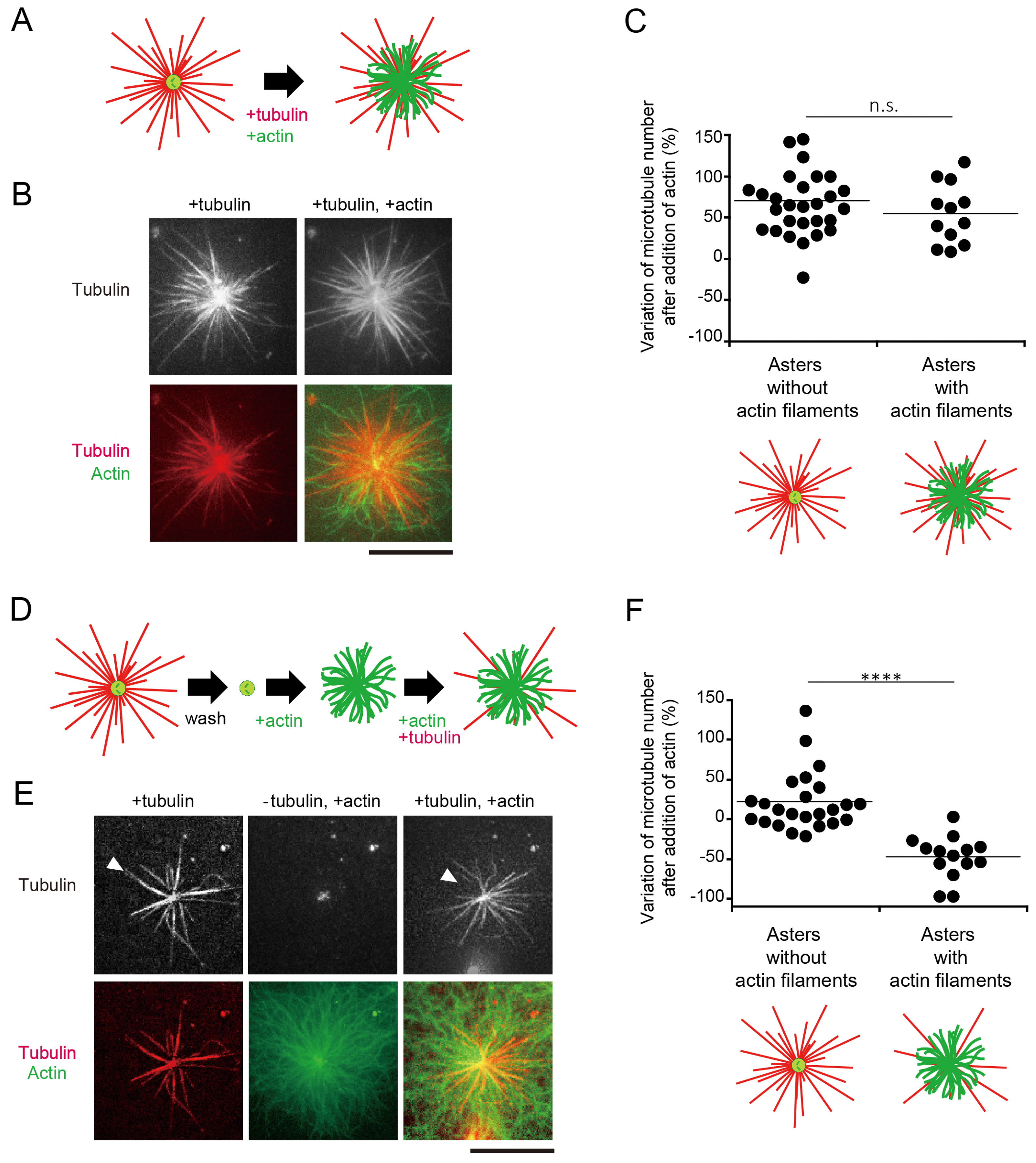
Steric hindrance of microtubule growth by actin filaments on isolated centrosomes. A- Schematic illustration of the first dynamic assay: sequential addition of tubulin followed by tubulin and actin on isolated centrosomes. B- Representative images showing microtubules (top line) and the merged images of actin and microtubules (bottom line) for the two steps of the assay; in the presence of tubulin only (left column) and in the presence of tubulin and actin (right column). Scale bar represents 10 μm. C- Quantification of the differences in the number of microtubules per centrosome between the two stages of the experiment described above on centrosomes capable (first condition), or not (second condition), to grow actin filaments. Data were collected from a single experiment. D- Schematic illustration of the second dynamic assay: tubulin is added to measure centrosome nucleation capacity and washed out. Then actin is added followed by actin and tubulin. E- Representative images showing microtubules (top line) and the merged images of actin and microtubule (bottom line) during the three steps of the assay; in the presence of tubulin only (left column), in the absence of tubulin and presence of actin (middle column) and in the presence of tubulin and actin (right column). Scale bar represents 10 μm. F- Quantification of the differences in the number of microtubules per centrosome between the first and last steps of the experiment described above (panels D and E) on centrosomes capable (first condition), or not (second condition), to grow actin filaments. Data were pooled from 2 independent experiments.

In a second experiment, tubulin dimers were initially added to quantify the number of microtubules per centrosome and in effect, to select those centrosomes with the capability to nucleate microtubules. The tubulin dimers and microtubules were then removed by rinsing the centrosomes in buffer. Actin monomers were then added, followed by tubulin dimers again (Figure 4D, E and F). For those centrosomes devoid of actin filaments, the microtubule number was not significantly different between the initial and final stages of the experiment (Figure 4F). By contrast, for centrosomes which nucleated actin filaments, the microtubule number was significantly reduced at the final stage than at the initial stage, confirming that actin filaments perturbed microtubule regrowth (Figure 4F).

### Microtubule growth is perturbed by actin filaments via steric hindrance in a biochemical model

In the above in vitro model, only 25% of the isolated centrosomes had the capability of nucleating microtubules, reflecting the difficulties in centrosome purification. Despite the optimisation steps to improve the quality of the centriole (Gogendeau et al., 2015), the isolation step results in centrosome with more or less fragmented peri-centriolar material. As a consequence, the investigation of their nucleation capacities was informative but intrinsically biased. Therefore, to directly test steric competition between actin and microtubules during the first stages of microtubule growth, we combined two distinct biochemical assays in which short microtubule seeds and actin nucleators were grafted onto the same microfabricated spot on a planar surface in vitro (Portran et al., 2013; Reymann et al., 2010) (Figure 5A).

**Figure 5:**
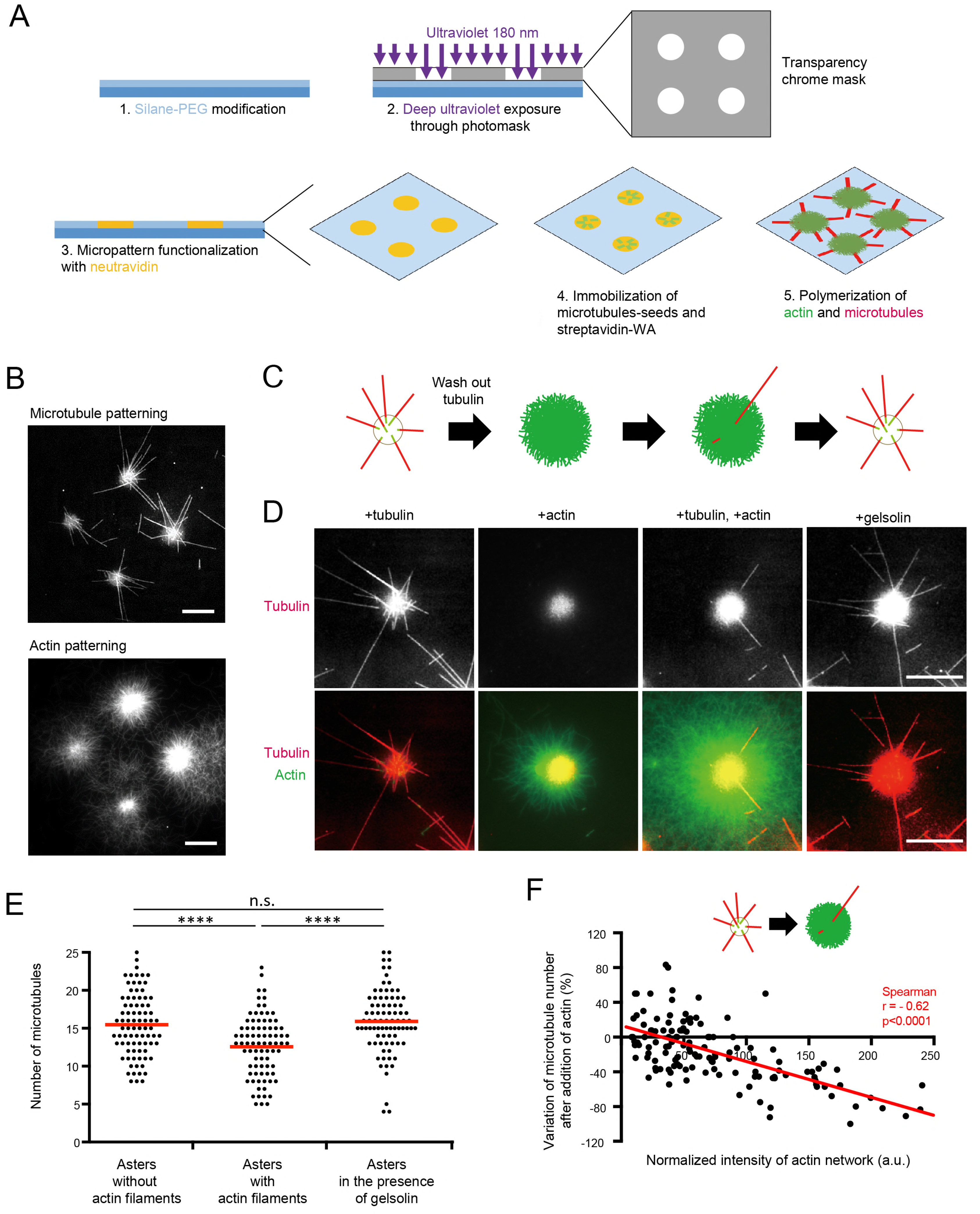
Reconstitution of the interplay between actin and microtubules on micropatterns. **A**- Schematic illustration of the micropatterning method to graft microtubules seeds (red) and actin-nucleation-promoting complexes (orange) on 8-micron-wide discoidal micropatterns. A glass coverslip (deep blue) coated with polyethyleneglycol (PEG) (light blue) was placed in contact with a transparency mask and exposed to deep UV light. The exposed coverslip was then immersed with neutravidin to fix biotinylated microtubule seeds (red) on exposed regions. Streptavidin-WA was immobilized on microtubule seeds via their interaction with biotin. Tubulin dimers and actin monomers were then added to allow filaments elongation. **B**- Representative images of microtubules (top) and actin filaments (bottom) growth from micropatterns. Scale bars represent 20 μm. **C**- Schematic illustration of the assay on micropatterned substrate. Tubulin was first added alone to measure the nucleation capacity of each micropattern, and then washed out. Later on, actin was added followed by actin and tubulin. Finally, actin was rinsed out and gelsolin was added to fully disassemble actin filaments. **D**- Representative images showing microtubules (top line) and the merged images of actin and microtubules (bottom line) during the four steps of the assay; in the presence of tubulin only, in the absence of tubulin and presence of actin, in the presence of tubulin and actin, and finally in the presence of tubulin and gelsolin but in the absence of actin filaments (from left to right). Scale bar represents 10 μm. **E**- Quantification of the number of microtubules per micropattern in the presence of tubulin only (left), actin and tubulin (middle) and tubulin only after actin filaments disassembly (right). Data were pooled from 2 independent experiments. **F**- The graph shows the same measurements as in panel E in an XY representation of individual measurements. It illustrates the differences in the number of microtubules per micropattern between the first to the second step (tubulin only versus actin and tubulin together) with respect to the density of actin filaments per micropattern.

In the biochemical model, the addition of free tubulin dimers and actin monomers led to the growth of both actin filaments and microtubules from each micropattern (Figure 5B). As with the in vitro model above, the micropattern were treated according to the following sequence: tubulin-dimer incubation, microtubule count; wash; actin-monomer incubation; and tubulin-dimer incubation (Figure 5C). The model showed again that microtubule formation was perturbed by the presence of actin filaments (Figure 5D, E). Interestingly, the addition of gelsolin to promote the disassembly of the actin filaments overcame the perturbation, indicating that the nucleation of actin filaments did not detach microtubule seeds but blocked their elongation (Figure 5C-E). Moreover, the relative density of actin was negatively correlated with microtubule numbers (Figure 5F). Therefore, given the absence of signalling pathways or cross-linking proteins, the perturbation of microtubule growth by actin filaments was by steric hindrance, and the denser the actin network, the greater the steric hindrance.

### Actin filament density at the centrosome is negatively affected by the degree of cell spreading

The experiments above supported the model in which at the centrosome, actin filaments perturb the formation of microtubules through steric hindrance. This led us to investigate how actin density at the centrosome is regulated in living cells. We have previously shown that with B-lymphocyte forming an immune synapse with antigen presenting cells actin nucleation is decreased at the centrosome (Obino et al., 2016). Because immune synapses are enriched for actin and adhesion molecules such as integrins, we hypothesized that the actin-filament density at the centrosome is inversely related to the degree of cell adhesion and spreading because actin nucleating structures compete for available actin monomers in the cell (Suarez and Kovar, 2016), Hence, minimal cell spreading permits a high amount of actin filaments to form at the centrosome thus perturbing microtubule growth, whereas extensive cell spreading sequesters most of the available actin monomers, reducing the number of actin filaments at the centrosome and thus favouring microtubule growth (Figure 6A).

**Figure 6:**
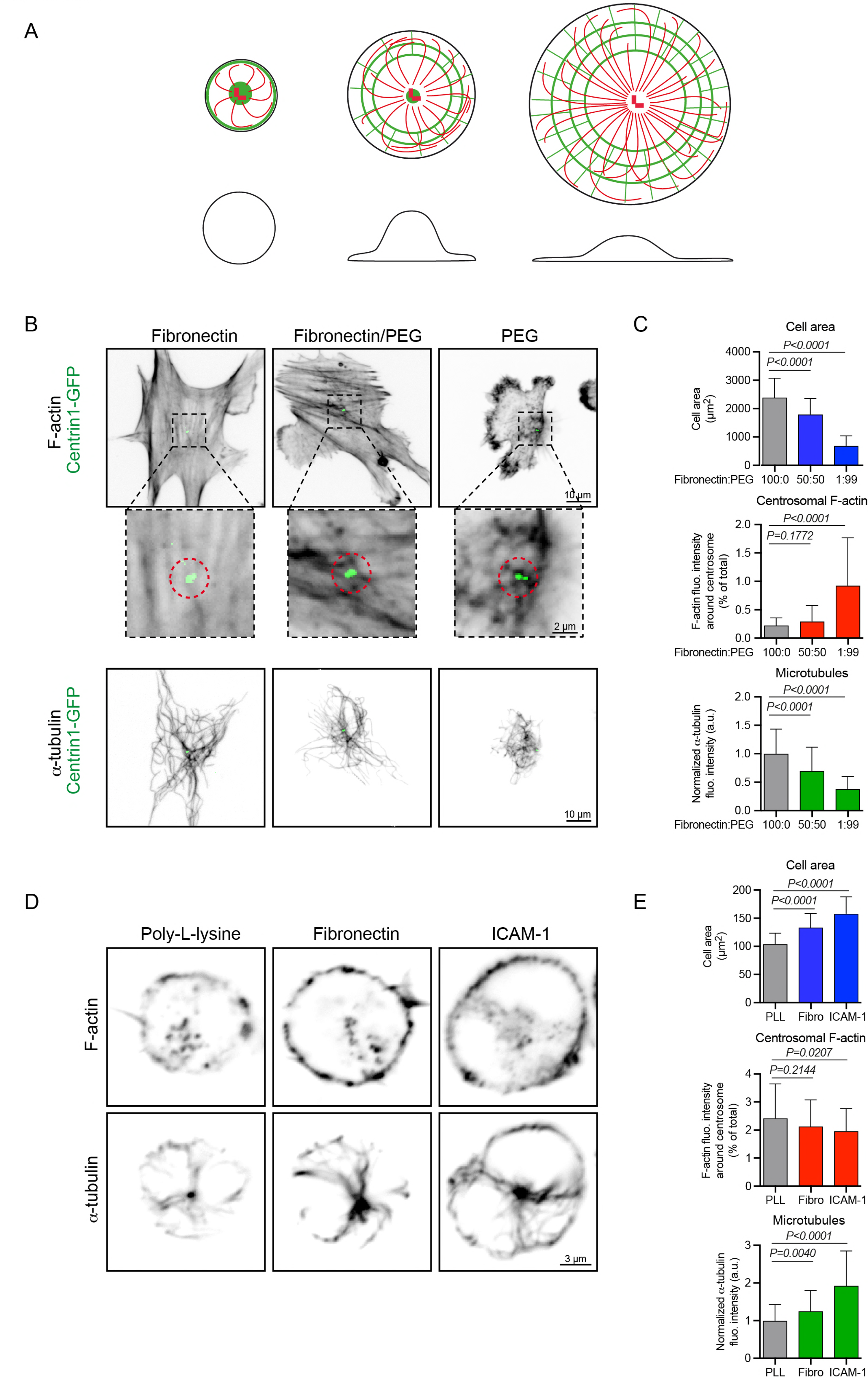
Modulation of microtubule growth by cell spreading and centrosomal actin. **A**- Schematic illustration of our model according to which cell spreading sequesters monomeric actin to the cortex and thereby enables the centrosome to grow more microtubules. Drawings show top (top line) and side views (bottom line) of cells with increased spreading from left to right. Actin filaments are in green, microtubules in red. **B**- RPE1 cells stably expressing centrin1-GFP were plated for 3 h on coverslips coated with different ratios (100:0; 50:50 or 1:99) of fibronectin and PLL-PEG prior to fixation and staining for F-actin (top line and magnified views around centrosome below. Scale bars represent 10 μm and 2 μm, respectively) and α-tubulin (bottom line. Scale bar represents 10 μm). **C**- Quantification of the area occupied by the cell on the substrate (top), centrosomal F-actin content (middle) and total amount of polymerized tubulin (bottom) for the three conditions of cell adhesion described in B. Measurements came from 3 independent experiments with more than 60 analyzed cells in each. Errors bars represent standard deviations. **D**- IIA1.6 B lymphoma cells were plated for 60 min on poly-L-lysine, fibronectin or ICAM-1 coated cover slides prior to be fixed and stained for F-actin (top line) and α-tubulin (bottom line). Scale bar represents 3 μm. **E**- Quantification of the area occupied by the cell on the substrate (top), centrosomal F-actin content (middle) and total amount of polymerized tubulin (bottom) for the three conditions of cell adhesion described in D. Measurements came from 3 independent experiments with more than 80 analyzed cells in each. Errors bars represent standard deviations.

For highly adherent RPE1 cells, three states of cell spreading (low, medium and high) were dictated by the degree of substrate adhesiveness (by tuning fibronectin concentration in PEG; Figure 6B). For low-adherent B-lymphocytes, three states of cell adhesion and spreading were dictated by plating on poly-lysine, fibronectin and ICAM-1 (Carrasco et al., 2004)(Figure 6C). For both cell types, the degree of cell adhesion and/or spreading (i.e. the area occupied by the cell on the substrate) was negatively correlated with centrosomal-actin density and positively correlated with the density of the microtubule network throughout the cell (Figure 6D, E). Altogether these results support a model in which microtubule growth from the centrosome is modulated by the adhesion state of the cell via the degree to which actin filaments are prevented from forming at the centrosome.

## Discussion

Actin is the most abundant protein in the cytoplasm and as such has long been considered as a major contaminant of centrosome proteomics studies (Bornens and Moudjou, 1999; Andersen et al., 2003). However, actin filaments have been directly observed at the poles of mitotic spindles (Stevenson et al., 2001; Chodagam et al., 2005) and at the centrosome of several cell types in interphase (Farina et al., 2016; Obino et al., 2016; Au et al., 2016). Centrosomal-actin filaments have been shown to anchor the centrosome the nucleus (Bornens, 1977; Burakov and Nadezhdina, 2013; Obino et al., 2016), support the transport of vesicles during ciliogenesis (Assis et al., 2017; Wu et al., 2018), connect basal bodies to the actin cortex in ciliated cells (Antoniades et al., 2014; Pan et al., 2007; Walentek et al., 2016) and power centrosome splitting in prophase (Uzbekov et al., 2002; Wang et al., 2008; Au et al., 2016).

The results of our study identify a new function for actin filaments at the centrosome. We propose a model in which these centrosomal actin filaments provide a conduit through which changes to actin networks at the cell periphery modulate the formation and growth of microtubules emanating from the centrosome. The centrosomal-actin filaments primarily perturb the formation of microtubules through steric hindrance. Indeed, physical constraints imposed by actin filaments (Huber et al., 2015) have been shown to limit microtubule shape fluctuations (Katrukha et al., 2017; Brangwynne et al., 2006) and centrosome displacement (Piel et al., 2000). However, we could not distinguish whether the actin filaments impaired nucleation or perturbed elongation. Moreover and in the cell, the assembly of actin filaments at the centrosome may expulse some microtubule nucleating factors or interfere with specific signalling pathways, in addition to sterically hindering microtubule growth.

Our results expand the description of cytoskeleton changes during B-lymphocyte activation (Obino et al., 2016) and show that centrosomal-actin filament disassembly promotes the growth of microtubules. Interestingly, the increase in microtubules may contribute to B cell polarization, a hallmark of their activation (Yuseff et al., 2011), by promoting centrosome off-centring. Indeed, a high quantity of microtubules can break network symmetry and force centrosome off-centring and its displacement to the cell periphery through the reorientation of pushing forces produced at the centrosome by microtubule growth (Letort et al., 2016; Burute et al., 2017; Pitaval et al., 2016). Therefore, centrosomal-actin filament disassembly could be involved in both the disengagement of the centrosome from the nucleus (Obino et al., 2016) and in the stimulation and reorganisation of microtubule-based pushing forces to drive centrosome motion toward the cell periphery.

The regulation of microtubule growth at the cell centre complements those mechanisms that regulate microtubule stability at the cell periphery, where microtubule stability is promoted by cell adhesions and their associated actin networks (Akhmanova and Steinmetz, 2015). Hence those mechanisms ensure a localised form of regulation that can directly bias microtubule network organisation (Gundersen et al., 2004; Etienne-Manneville, 2013). At the cell centre, microtubule growth adaptation to cell shape, cell adhesion and cell spreading can be mediated by the centrosomal-actin network (Figure 6A). An explanation for this is that cell adhesion and spreading triggers the elaboration of actin networks at the cortex, hence reducing the pool of available actin monomers, and potentially sequestering from the centrosome actin-filament nucleation and branching factors such as Arp2/3 and WASH (Suarez and Kovar, 2016) (Obino et al., 2016)(Farina et al., 2016). The reduction in the centrosomal-actin network thus allows more microtubules to be nucleated at the centrosome. The interplay at the centrosome between actin filaments and microtubules in response to cell spreading may have important implications for the ability of the cell to sense and adapt to external cues.

## Materials and Methods

### Cell culture

Stable Jurkat cell lines expressing centrin1-GFP (Farina et al., 2016) were cultured in RPMI 1640, (Gibco). Cells were not sorted based on GFP fluorescence. The mouse B lymphoma cell line IIA1.6 (derived from the A20 cell line (American Type Culture Collection #: TIB-208)) was cultured as reported (Obino et al., 2016) in CLICK medium (RPMI1640—GlutaMax-I), supplemented with 0.1% β-mercaptoethanol and 2% sodium pyruvate. The RPE1 cell line stably expressing centrin1-GFP (Farina et al., 2016) was cultured in DMEM/F-12. All media were supplemented with 10% fetal calf serum and penicillin/streptomycin. Cells were cultured at 37°C and 5% CO_2_. All cell lines were tested monthly for mycoplasma contamination.

Cytoskeleton inhibitors (CK666 at 25 μM, SMIFH2 at 25μM; Latrunculin-A at 5μM; all from Tocris Bioscience) were added in the cell medium for 45 minutes at 37 °C.

For the coating of glass coverslips; fibronectin (Sigma Aldrich) was used at 10 μg/ml and PLL-PEG (JenKem Technologies, Texas) at 10 μg/ml in Hepes 10 μM, Poly-L-Lysine (Invitrogen) was used at 10 μg/mL, and ICAM-1 (R&D System) was used at 10 μg/mL.

### Preparation of BCR-ligand-coated beads

Latex NH_2_-beads 3 μm in diameter (Polyscience) were coated with 8% glutaraldehyde (Sigma Aldrich) for 2 h at room temperature (4×10^7^ beads/ml). Beads were washed with PBS and incubated overnight at 4°C with 100 μg/ml of either F(ab’)_2_ goat anti-mouse IgG (BCR-ligand^+^ beads) or F(ab’)_2_ goat anti-mouse IgM (BCR-ligand^−^ beads; MP Biomedical).

### Cell fixation and immuno-staining

Cells were extracted by incubation for 15 sec with cold Cytoskeleton buffer (10 mM MES pH6.1, 138 mM KCl, 3 mM MgCl_2_, 2 mM EGTA) supplemented with 0.5% Triton X-100 and fixed with Cytoskeleton buffer supplemented with 0.5% glutaraldehyde for 10 min at room temperature. Glutaraldehyde was reduced with 0.1% sodium borohydride (NaBH_4_) in 1x PBS for 7 min and unspecific binding sites were saturated using a solution of 1x PBS supplemented with 2% BSA and 0.1% Triton X-100 for 10 min. The following primary antibodies were used: monoclonal rat anti-α-tubulin (AbD Serotec, Clone YL1/2, 1/1000) and human anti-green fluorescent protein (GFP) (Recombinant Antibody Platform, Institut Curie, Paris, France, 1/200). The following secondary antibodies were used: AlexaFluor647-conjugated F(ab’)_2_ donkey anti-rat and AlexaFluor488-conjugated donkey anti-human (Life Technologies, both 1/200). F-actin was stained using AlexaFluor546-conjugated phalloidin (Life Technologies, #A22283, 1/100).

### Isolation of centrosomes

Centrosomes were isolated from Jurkat cells by modifying a previously published protocol (Moudjou and Bornens, 1998; Gogendeau et al., 2015). In brief, cells were treated with nocodazole (0.2 μM) and cytochalasin D (1 μg/ml) followed by hypotonic lysis. Centrosomes were collected by centrifugation onto a 60% sucrose cushion and further purified by centrifugation through a discontinuous (70%, 50% and 40%) sucrose gradient. The composition of the sucrose solutions was based on a TicTac buffer, in which the activity of tubulin, actin and actin-binding proteins is maintained: 10 mM Hepes, 16 mM Pipes (pH 6.8), 50 mM KCl, 5 mM MgCl_2_, 1 mM EGTA. The TicTac buffer was supplemented with 0.1% Triton X-100 and 0.1% β-mercaptoethanol. After centrifugation on the sucrose gradient, supernatant was removed until only about 5 ml remained in the bottom of the tube. Centrosomes were stored at −80 °C after flash freezing in liquid nitrogen.

### Protein expression and purification

Tubulin was purified from fresh bovine brain by three cycles of temperature-dependent assembly/disassembly in Brinkley Buffer 80 (BRB80 buffer: 80 mM Pipes pH 6.8, 1 mM EGTA and 1 mM MgCl_2_) (Shelanski, 1973). Fluorescently labelled tubulins (ATTO-488 and ATTO-565-labelled tubulin) were prepared by following previously published method (Hyman et al., 1991).

Actin was purified from rabbit skeletal-muscle acetone powder. Monomeric Ca-ATP-actin was purified by gel-filtration chromatography on Sephacryl S-300 at 4°C in G buffer (2 mM Tris-HCl, pH 8.0, 0.2 mM ATP, 0.1 mM CaCl_2_, 1 mM NaN_3_ and 0.5 mM dithiothreitol (DTT)). Actin was labelled on lysines with Alexa-488 and Alexa-568. Recombinant human profilin, mouse capping protein, the Arp2/3 complex and GST-streptavidin-WA were purified in accordance with previous methods (Michelot et al., 2007; Achard et al., 2010).

### In vitro assays with isolated centrosomes

Experiments were performed in polydimethylsiloxane (PDMS) stencils in order to add/exchange sequentially experimental solutions when needed. PDMS (Sylgard 184 kit, Dow Corning) was mixed with the curing agent (10: 1 ratio), degassed, poured into a Petri dish to a thickness of 5 mm and cured for 2 h at 80 °C on a hot plate. The PDMS layer was cut to square shape with dimension of 10 mm × 10 mm and punched using a hole puncher (Ted Pella) with an outer diameter of 6 mm. The PDMS chamber were oxidized in an oxygen plasma cleaner for 40 s at 60W (Femto, Diener Electronic) and brought it into contact with clean coverslip (24 mm × 30 mm) via a double sided tape with 6 mm hole.

Isolated centrosomes were diluted in TicTac buffer (Farina et al., 2016) and incubated for 20 min. To remove excess of centrosomes and coating the surface of coverslips, TicTac buffer supplemented with 1% BSA was perfused into PDMS chamber, which was followed by a second rinsing step with TicTac buffer supplemented with 0.2% BSA and 0.25% w/v methylcellulose. Microtubules and actin assembly at the centrosome were induced using a reaction mixture containing tubulin dimers (labelled with ATTO-488 or ATTO-565, 18 μM final) and actin monomers (labelled with Alexa-488 or Alexa-568, 0.3 – 1.0 μM final) in TicTac buffer supplemented with 1 mM GTP and 2.7 mM ATP, 10 mM DTT, 20 μg/ml catalase, 3 mg/ml glucose, 100 μg/ml glucose oxidase and 0.25% w/v methylcellulose. In addition, a threefold molar equivalent of profilin to actin and 60 nM Arp2/3 complex were added in the reaction mixture.

Sequential microtubule and actin assembly experiments were carried out based on the aforementioned method. In brief, after assembling microtubules by adding tubulin in the reaction mixture (18 μM final) for 15 min, microtubules were removed by exchanging the reaction mixture with TicTac buffer supplemented with 0.2% BSA and 0.25% w/v methylcellulose. Subsequently, the reaction mixture of actin (1 μM final) with profilin and Arp2/3 was applied to assemble the actin aster. After 15 min incubation, the tubulin reaction mixture with actin, profilin and Arp2/3 complex was added to assemble both microtubules and actin asters together.

### Micropatterning

Micropatterning of microtubules and actin filaments were performed in accordance with previous published methods with modification (Reymann et al., 2010; Portran et al., 2013). In brief, cleaned glass coverslips were oxidized with oxygen plasma (5 min, 60 W, Femto, Diener Electronic) and incubated with poly-ethylene glycol silane (5 kDa, PLS-2011, Creative PEGWorks, 1 mg/ml in ethanol 96.5% and 0.02% of HCl) solution for overnight incubation. PEGylated coverslips were placed on a chromium quartz photomask (Toppan Photomasks, Corbeil, France) using a vacuum holder. The mask-covered coverslips were then exposed to deep ultraviolet light (180 nm, UVO Cleaner, Jelight Company, Irvine, CA) for 5 min. The PDMS open chamber was assembled as described above. Neutravidin (0.2 mg/ml in 1 × HKEM [10 mM Hepes pH 7.5, 50 mM KCl, 5 mM MgCl_2_, 1 mM EGTA]) was perfused in PDMS chamber and incubated for 15 min. The biotinylated microtubule seeds, which were prepared with 20% of fluorescent-dye-labelled tubulin and 80% biotinylated tubulin in presence of 0.5 mM of GMPCPP as previously described (Portran et al., 2013), were deposited on neutravidin-coated surface. Subsequently, 1 μM of streptavidin-WA in 1x HKEM was added into the PDMS chamber. After each step, the excess of unbound proteins was washed away using wash buffer. Microtubules and actin filaments were assembled according to the above protocol (see *In vitro assays*), except that 120 nM of Arp2/3 complex was used instead of 60 nM. To disassemble actin asters on the micropatterns, gelsolin (1.6 μM, gift from Robert Robinson laboratory, IMCB Singapore) was added into the reaction mixture at the last step of the experiment.

### Imaging and analysis

All z-stack images (0.5 μm spacing) of fixed cells were acquired on an inverted spinning disk confocal microscope (Roper/Nikon) with a 60x/1.4 NA oil immersion objective. Image processing was performed with Fiji (ImageJ) software. Centrosome-associated F-actin was quantified as previously described (Obino et al., 2016). Briefly, after selecting manually the centrosome plane, we performed a background subtraction (rolling ball 50 px) on the z-projection (by calculation of pixel average intensity) of the three planes above and below the centrosome. The total fluorescence of centrosomal F-actin was measured in a 1.6-micrometers-wide disc centred on the centrosome and the total fluorescence of microtubule was measured in the entire cell.

The imaging of microtubules, actin filaments and centrosomes in the in vitro experiments was performed with a total internal reflection fluorescence (TIRF) microscope (Roper Scientific) equipped with an iLasPulsed system and an Evolve camera (EMCCD) using using 60x Nikon Apo TIRF oil-immersion objective lens (N.A=1.49). The microscope stage was maintained at 37 °C by means of a temperature controller to obtain an optimal microtubule growth. Multi-stage time-lapse movies were acquired using Metamorph software (version 7.7.5, Universal Imaging). Actin-nucleation activity was quantified by measuring the actin-fluorescence intensity integrated over a 20 μm diameter at the centre of the actin-aster and normalized with respect to initial background intensity. The number of microtubules was manually counted from fluorescence microscopy images. All the measurements were done using Adobe Photoshop CC and the corresponding graphs were produced using Kareidagraph 4.0.

### Statistics

For the in vitro experiments (Figure 3, 4 and 5), statistical differences were identified using the unpaired *t*-test with Welch’s correction and Kaleidagraph software. For the cellular studies (Figure 1, 2 and 6) statistical differences were computed using GraphPad Prism 7 Software. No statistical method was used to determine sample size. Kolmogorov–Smirnov test was used to assess normality of all data sets. The following tests were used to determine statistical significance: Figures 1B, 2B, 2F, 3D, 4C, 6C (actin and microtubules) and 6E (actin and microtubules): Mann–Whitney test; Figures 4A, 5A, 6C (cell area) and 6E (cell area): Unpaired t test; Figures 1C, 2C and 2G: One sample t test (comparison to a theoretical mean of zero, where zero represents no difference between conditions); Figures 1D, 2D, 2H and 5E: Spearman correlation test. ^****^P < 0.0001, ^***^P < 0.001, ^**^P < 0.01, ^*^P < 0.05, n.s=not significant. Bar graphs describe the mean ± standard deviation.

## Acknowledgements

We are grateful to the CytoMorpho Lab for stimulating discussions and the collective effort to perform the experiments.

## Author contributions

DI performed all experiments in reconstituted systems. DO performed all experiments in living cells. FF performed preliminary experiments in reconstituted systems. JG and CG were involved in experiments in reconstituted systems. A-ML-D, LB and MT designed the project, obtained the funding to support it and supervised the work. MT wrote the manuscript. All authors reviewed/edited the manuscript for important intellectual content.

